# Quantitative MRI Reveals Bone Marrow Regeneration Following Targeted Marrow Irradiation and Transplantation in a Sickle Cell Disease Model

**DOI:** 10.64898/2025.12.08.692634

**Authors:** Malakeh Malekzadeh, Hemendra Ghimire, Ji Eun Lim, Srideshikan S Madabushi, Weidong Hu, Marco Andrea Zampini, Kazuki Fujita, Raghda Fouda, Guy Storme, Susanta K. Hui

## Abstract

Sickle cell disease (SCD) is associated with chronic bone marrow stress, altered hematopoiesis, and reduced adiposity. Whether marrow-selective conditioning followed by transplantation normalizes these abnormalities remains unclear. We investigated bone marrow remodeling in Townes mice by comparing SCD control (SCD-Con) with mice that received total marrow irradiation (TMI) followed by bone marrow transplantation (SCD-TMI-BMT). Multiparametric micro-MRI at 7 T quantified proton density water fraction (PDWF), proton density fat fraction (PDFF), and R2*(1/T2*), and micro-CT assessed trabecular structure in the femur.

SCD-Con marrow showed higher water content (elevated PDWF), reduced adiposity (lower PDFF), and imaging features consistent with erythroid hyperplasia and elevated iron burden (shorter T2* with reciprocal increase in R2*). In contrast, SCD-TMI-BMT mice demonstrated smaller R2*, reduced PDWF, and partial restoration of adiposity, accompanied by reciprocal shifts in R2*, consistent with decreased cellular iron and marrow remodeling. Micro-CT revealed an improved trabecular architecture after BMT compared to SCD control. MRI imaging biomarkers aligned with histologic evidence of reduced cellularity and larger adipocyte voids.

In conclusion, a TMI-BMT SCD model promotes partial normalization of the marrow microenvironment. Multiparametric MRI with micro-CT provides a practical, non-invasive framework for monitoring marrow remodeling and skeletal health after curative therapy.

## 1. Introduction

Sickle cell disease (SCD) is the most common inherited blood disorder in the United States, impacting around 100,000 individuals nationally and leading to approximately 300,000 new cases worldwide each year(^1, 2^). SCD is caused by a single point mutation in the beta-globin gene, substituting valine for glutamic acid at the sixth position, resulting in abnormal hemoglobin (HbS)(^3^). While the condition typically arises from a homozygous HbSS genotype, it can also occur in combinations such as HbS with Hemoglobin C (HbC) or other hemoglobinopathies, like beta-thalassemia (e.g., HbSβ0). The disease’s core pathology involves polymerization of deoxygenated HbS, distorting red blood cells from their usual biconcave shape into rigid, fragile sickle-shaped cells(^4^). Bone marrow transplantation (BMT) is one of the few curative therapies for SCD, requiring conditioning to remove native hematopoietic stem cells that carry the β-globin S mutation(^5^). Traditional conditioning using total body irradiation (TBI) for hematopoietic cell transplantation is risky for SCD patients due to high toxicity and organ damage. However, the associated toxicities of high-dose TBI often outweigh its survival benefits. To address this limitation, we developed CT-based, image-guided TMI platforms(^6, 7^) that concentrate dose in marrow while sparing critical organs (gastrointestinal tract, lungs, heart, liver) and support durable donor engraftment in immunocompetent wild-type mice. Compared with TBI, TMI significantly reduced acute and chronic organ injury(^8^) and enhanced repair, reflected by greater villus height and a lower areg/egf mRNA ratio. Organ sparing with TMI did not compromise engraftment, reduce normal tissue toxicity and supports dose-escalation studies even in the aging mouse model.

In a SCD mouse model, TMI–based conditioning with BMT improves donor engraftment and systemic hematologic indices(^9^), but its effects on the marrow microenvironment and bone quality remain undefined. Conventional assessments rely on terminal histology and limited sampling, which precludes longitudinal evaluation. To address this gap, we used quantitative MRI to map marrow composition through proton-density water and fat fractions (PDWF, PDFF) and to quantify susceptibility-driven relaxation metrics T2* and R2* as proxies for cellular and iron burden, and we used micro-CT to quantify femoral trabecular architecture. This noninvasive, multimodal approach tests whether TMI-BMT promotes marrow remodeling and skeletal improvement and sets up the quantitative MRI and micro-CT measurements described next to interrogate marrow composition, iron burden, and trabecular structure.

Quantitative MRI offers noninvasive advantages over blood tests, biopsies, and qualitative imaging for assessing marrow health, iron burden, and treatment response. Although in-vivo spatial mapping has technical constraints, advanced methods such as water-fat separation and relaxation mapping can detect subtle skeletal changes. Marrow content can be quantified using PDFF and PDWF derived from water-fat chemical-shift separation(^10–13^). Moreover, T2* reflects the combined effects of spin–spin interactions and microscopic field inhomogeneities, whereas R2* (1/T2*) serves as a quantitative marker of susceptibility effects and is widely used for iron assessment(^11, 12^). SCD is associated with skeletal fragility, characterized by low bone mineral density and impaired trabecular microarchitecture(^14^). Thus, combining MRI with micro-CT enables simultaneous evaluation of compositional and architectural remodeling after TMI-BMT.

Because TMI is image-guided radiotherapy, pairing it with quantitative MRI and micro-CT enables noninvasive, in vivo tracking of marrow depletion and regeneration and the accompanying skeletal remodeling, reducing reliance on terminal pathology and informing treatment monitoring. In this study, the primary endpoint was quantitative MRI of the right femur (PDWF, PDFF, T2*, R2*) to assess marrow composition and susceptibility. Secondary endpoints quantified trabecular microarchitecture by high-resolution micro-CT and tertiary endpoints included H&E histology for qualitative concordance of cellularity and adipocyte morphology. Together, these modalities allow simultaneous evaluation of compositional and architectural remodeling after TMI-BMT.

## 2. Methods

### 2.1. Study Design and TMI Treatment

This cross-sectional, non-randomized, unblinded preclinical study included 18 Townes SS mice allocated to two groups: SCD-TMI-BMT (n=8) and age- and sex-matched SCD controls (SCD-Con; n=10). At 6 months of age, SCD-TMI-BMT mice received CT-guided TMI (10 Gy) in which 10 Gy to marrow and 4 Gy to the remainder of the body, delivered as in two fractions of 5 Gy separated by 6 hours, followed 24 hours later by intravenous transplantation of 25×10^6^ whole bone-marrow cells from Townes AA donors (10–12 weeks, non-sickling). TMI was delivered on a Precision X-RAD SmART Plus/225cx system (Precision X-Ray, North Branford, CT, USA)(^9^). Imaging was performed at ∼4 months post-transplant (age ∼10 months): quantitative MRI first, micro-CT 3 days later, and histology 3 days after micro-CT (Fig. 1A). Donor chimerism and stem-cell marker analyses were reported previously(^9^) and were not repeated here.

**Figure 1.**
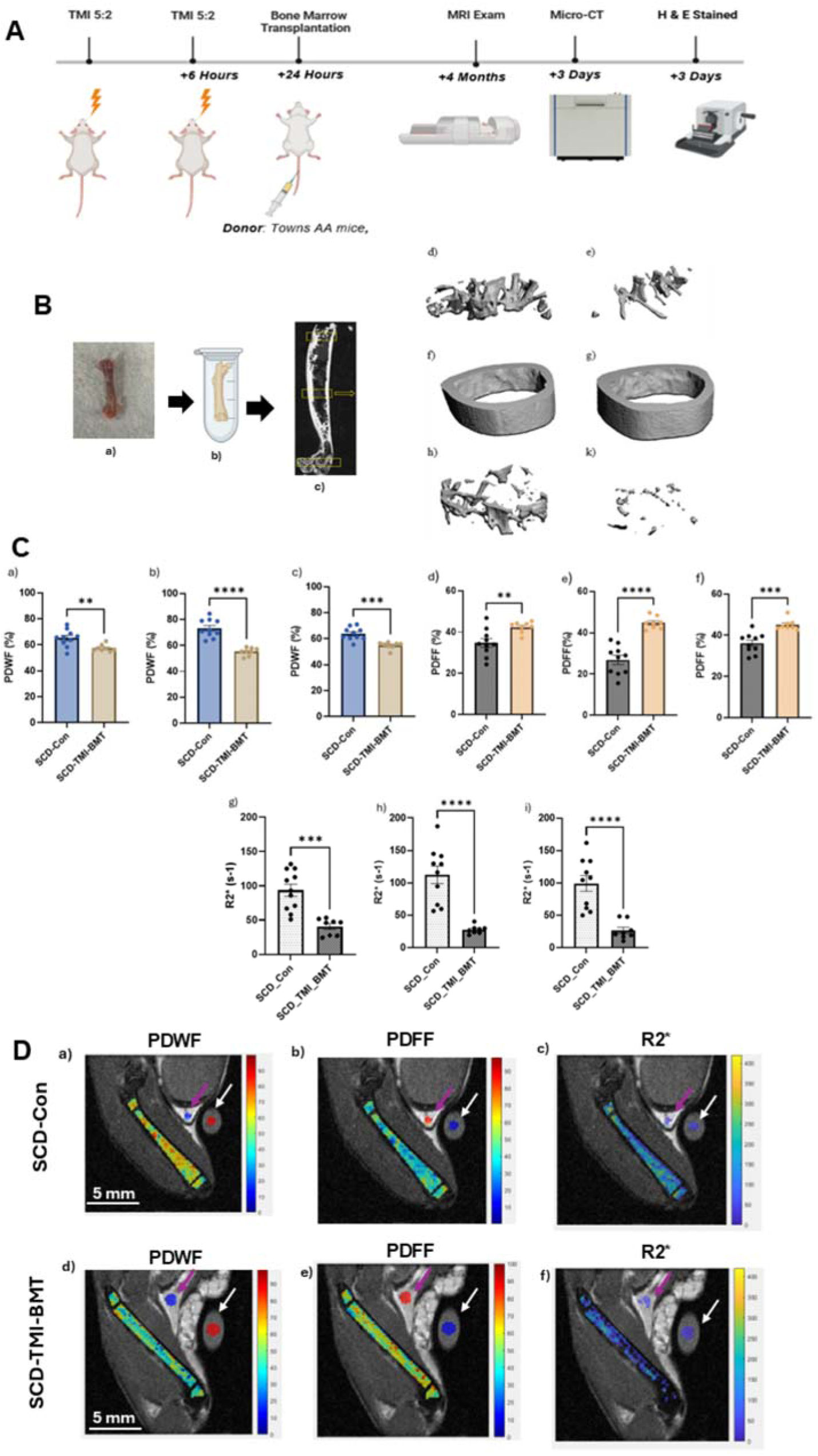
Experimental timeline and multimodal assessment of bone and marrow in SCD-Control and SCD-TMI-BMT mice. (A) Schematic of the experimental timeline. a) Six-month-old Townes SS mice underwent total marrow irradiation (TMI) in two doses (5 Gy followed by 2 Gy, 6 h apart), followed by 24h transplantation of 25 million whole bone marrow cells from healthy Townes AA donor mice. At 4 months post-transplantation (age ∼10 months), mice underwent MRI (right femur, supine position) to assess bone marrow composition. The same femur was collected for micro-CT imaging within 3 days and for H&E histological analysis 3 days later to evaluate bone structure and marrow morphology. (B) Representative micro-CT imaging workflow. a) Extracted femur sample; b) Placement of the sample in a microtube for scanning; c) Coronal micro-CT slice indicating anatomical landmarks used to define the proximal, mid-shaft, and distal regions of interest (ROIs), 3D micro-CT reconstructions highlighting the femoral regions in SCD-Control (d, f, h) and SCD-TMI-BMT (e, g, k) mice: proximal (d, e), mid-shaft cortical (f, g), and distal (h, k) views. (C) MRI-derived parameters for the proximal (a, d, and g,), mid-shaft (b, e, and h), and distal (c, f, and j) regions of the femur: PDWF (a-c), PDFF (d-f), and R2* (g-j). (D) PDWF, PDFF, and R2* maps map overlaid on the T2 image for three femoral ROIs (proximal, mid-shaft, distal), with additional circular ROIs for subcutaneous fat (purple arrow) and a water tube (white arrow). Representative examples are shown for PDWF map and (b) PDFF, and (c) R2* map for the SCD-Con mouse (a–c) and one SCD-TMI-BMT mouse (d–f). *PDWF, proton density water fraction; PDFF, proton density fat fraction; R2*, effective transverse relaxation rate; SCD-Con, SCD control; and SCD-TMI-BMT, SCD total marrow irradiation treated. Statistical differences were evaluated, and significance is denoted as follows: ns = not significant, *p < 0.05, **p < 0.01, ***p < 0.001, ****p < 0.0001, (Each dot represents one mouse). (MRI was done on a preclinical 7 T MRI-PET system (MR Solutions Ltd., UK)*.

### 2.2. Quantitative Magnetic Resonance Imaging

Imaging was performed on a 7 T MRI-PET system (MR Solutions Ltd., UK). Anesthetized mice were positioned supine with femurs aligned for coronal imaging; body temperature was maintained at 37 °C with respiratory gating. A water-filled tube served as an external signal reference (Fig.S1). For water–fat separation, a multi-echo spoiled gradient-echo (ME-GRE) scan was acquired over the right femur. Binary regions of interest (ROIs) were defined a priori at the proximal metaphysis, midshaft diaphysis, and distal metaphysis of the right femoral marrow. PDFF and PDWF maps were computed in MATLAB from fat- and water-only images as PDFF = Fat/ (Water + Fat) × 100 and PDWF = Water/ (Water + Fat) × 100.

Paramagnetic ferritin and hemosiderin cause local field inhomogeneities that shorten T2*(^10, 14^) where the effective relaxation rate is R2* = 1/T2*. T2* mapping used the Multi-echo gradient-recalled echo (ME-GRE) dataset (repetition time (TR)/echo time (TE)= 500 ms / 3.0, 5.5, 8.0 ms). Voxel-wise signal decay was fit with a monoexponentially model, 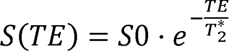, using nonlinear least squares; results are reported as R2* (s⁻¹), with T2* (ms) presented as 1/R2* for interpretability. Detailed sequence parameters, shimming procedure, coil configuration, preprocessing, ROI propagation, and quality control are provided in the Supplementary Methods.

### 2.3. Micro-CT

After MRI, right femurs were fixed and scanned ex-vivo on a μCT 50 (Scanco Medical, city, country) prior to decalcification and H&E (Fig.1B, a & b). Trabecular morphology and bone mineral density (BMD) were quantified in three predefined ROIs (proximal metaphysis, midshaft diaphysis, distal metaphysis) (Fig.1B, c). Acquisition parameters, Volume-of-interest (VOI) definitions, reconstruction/thresholding, and the full list of morphometric outputs are provided in the Supplementary Methods.

### 2.4. Hematoxylin and eosin staining (H&E)

Bone marrow cellularity was evaluated on H&E-stained femoral sections at 10× magnification. Details of the staining protocol are in the supplementary documentation.

### 2.5. Statistical analysis

Statistics were performed in GraphPad Prism (v10.3.1). Normality was tested (Shapiro–Wilk); Parametric data: t-tests, Pearson; nonparametric: Mann–Whitney U, Spearman. Two-way ANOVA assessed effects of Group, Region, and their interaction, with significance set at *p* < 0.05.

## 3. Results

### 3.1. MRI

SCD-TMI-BMT mice (n=8) were compared with SCD-Con mice (n=10). At four months post-TMI-BMT, quantitative MRI of the right femur showed significantly higher PDWF in SCD-Con compared to SCD-TMI-BMT across proximal, midshaft, and distal ROIs (Fig. 1C, a–c), with the inverse pattern for PDFF (Fig. 1C, d–f).

R2* values were lower in SCD-TMI-BMT in all ROIs (Fig. 1C, g-i), where SCD-Con exhibited higher PDWF and lower PDFF than SCD-TMI-BMT. The mean values with their 95% confidence intervals (CI) for all ROIs and groups are shown in Table 1.

**Table 1.** Quantitative MRI measurements of the proximal, mid-shaft, and distal femur in SCD-Con and SCD-TMI-BMT groups. Values are presented as mean ± 95% CI.

| Group | ROI | PDWF (%) | PDFF (%) | R2*(s<sup>-1</sup>) |
| --- | --- | --- | --- | --- |
| | | mean ± 95% CI | mean ± 95% CI | mean ± 95% CI |
| SCD-Con | Proximal | 65.23 ± 4.68 | 34.77 ± 4.68 | 93.57 ± 19.53 |
| | Mid-Shaft | 73.14 ± 5.01 | 26.87 ± 5.01 | 112.6 ± 30.58 |
| | Distal | 63.80 ± 3.48 | 36.20 ± 3.48 | 99.41 ± 27.71 |
| SCD-TMI-BMT | Proximal | 57.55 ± 2.22 | 42.46 ± 2.21 | 40.60 ± 10.07 |
| | Mid-Shaft | 55.22 ± 2.51 | 44.78 ± 2.51 | 27.68 ± 5.28 |
| | Distal | 54.90 ± 2.20 | 45.10 ± 2.20 | 26.68 ± 11.75 |

PDWF, proton density water fraction; PDFF, proton density fat fraction; R2\*, effective transverse relaxation rate; SCD-Con, SCD control; and SCD-TMI-BMT, SCD total marrow irradiation treated.

Representative PDWF, PDFF, and R2* maps are presented for three femoral ROIs (proximal, mid-shaft, and distal), together with circular ROIs for subcutaneous fat and a reference water tube. Examples are shown for one SCD-Con mouse (Fig. 1D, a-c) and one SCD-TMI-BMT mouse (Fig. 1D, d-f).

We observed a group effect for PDWF, PDFF, and R2*, but only R2* showed an effect independent of ROI (Fig.2A). Two-way ANOVA showed a significant group effect for PDWF (F(1,48) = 70.54, p < 0.0001; SCD-Con > SCD-TMI-BMT) and PDFF (F(1,48) = 24.88, p < 0.0001; SCD-TMI-BMT > SCD-Con), (Fig 2A. a, b). Both also showed significant ROI effects (PDWF: F(2,48) = 4.18, p = 0.021; PDFF: F(2,48) = 4.11, p = 0.021) and Group × ROI interactions (F(2,48) = 5.56, p = 0.0068), indicating regional variation. For R2*, a strong group effect was found ( F (1,48) = 80.24, p < 0.0001; SCD-TMI-BMT < SCD-Con), (Fig.2A.c), with no ROI (p = 0.749) or interaction effects (p = 0.326), confirming uniform treatment effects across regions.

**Figure 2.**
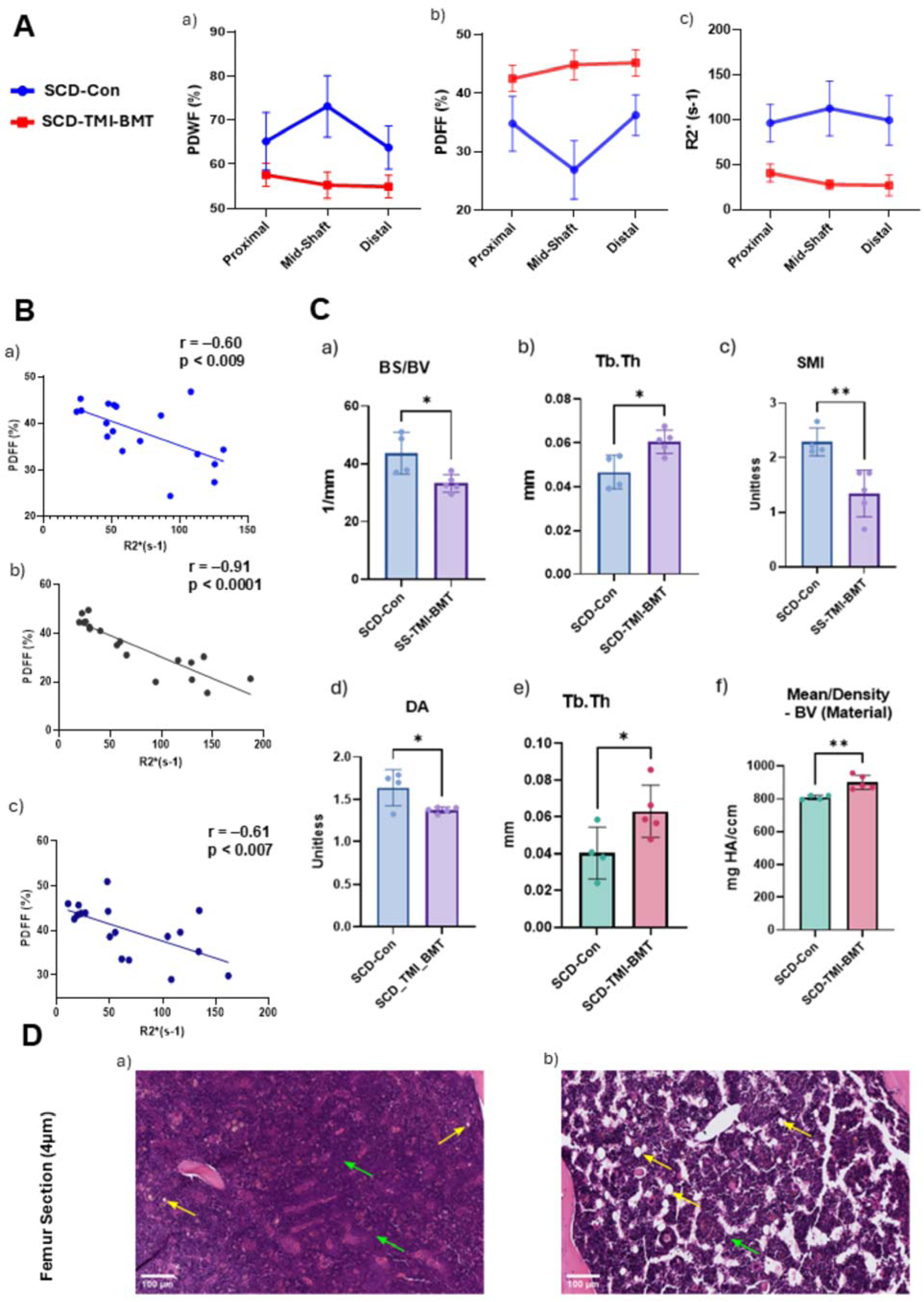
(A) Line graph depicting mean values with 95% confidence intervals across three ROIs in SCD-Con and SCD_TMI_BMT groups. (a) Two-way ANOVA revealed a significant main effect of group for PDWF (p < 0.0001), with SCD-Con showing higher values compared to SCD TMI. A significant effect of ROI was also observed (p = 0.021), along with a significant Group × ROI interaction (p = 0.0068), indicating that the treatment effect varied across ROIs. (b) Two-way ANOVA revealed a significant main effect of group for PDFF (*p* < 0.0001), with SCD-TMI-BMT showing higher values compared to SCD-Con. A significant effect of ROI was also observed (*p* = 0.021), along with a significant Group × ROI interaction (*p* = 0.0068), indicating that the treatment effect varied across ROIs. (c) Line graph (mean R2* ± CI95%) across three ROIs in SCD-Con and SCD-TMI-BMT groups (p <0.0001), with no ROI (p = 0.749) or interaction effects (p = 0.326). **(B) The correlation analysis between PDFF and R2*across three regions of interest (ROIs).** a) Proximal: Scatter plot with best-fit line showing the Pearson’s correlation between PDFF and R2* (rl1l = −0.598, p=0.0088). b) Mid-Shaft: Scatter plot with best-fit line showing the Spearman’s correlation between PDFF and R2* (rl1l = −0.913, p <0.0001). c) Distal: Scatter plot with best-fit line showing the Pearson’s correlation between PDFF and R2* (rl1l = −0.611, p <0.007). **(C) Micro-CT-based quantitative analysis of trabecular bone**, a) bone surface to volume ratio (BS/BV), b) trabecular thickness (Tb.Th), c) structure model index (SMI), d) degree of anisotropy (DA) in the proximal including; e) Tb.Th, and f) bone volume density (BV) distal femur including. **(D Histology**. Representative H&E-stained femoral marrow sections(4 µm) at 10x magnification showing restoration of marrow adipocytes (yellow arrows) and reduced erythroid hyperplasia (green arrows) in SCD-TMI-BMT mice compared with diffuse hypercellularity and adipocyte loss in SCD-Con. Images analyzed using QuPath. PDWF, proton density water fraction; PDFF, proton density fat fraction; R2*, effective transverse relaxation rate; SCD-Con, SCD control; and SCD-TMI-BMT, SCD total marrow irradiation treated. Statistical differences were evaluated, and significance is denoted as follows: ns = not significant, *p < 0.05, **p < 0.01, ***p < 0.001, ****p < 0.0001, (Each dot represents one mouse).

### 3.2. Association between R2* and PDFF

The correlation analysis demonstrated a significant relationship between marrow R2* and PDFF across all evaluated ROIs. When combining subjects from both groups (SCD-Con and SCD-TMI-BMT) within each ROI, a moderate-to-strong negative correlation was observed: Pearson’s correlation in the proximal region (r = –0.60, p < 0.009), (Fig.2B.a), Spearman’s correlation in the mid-shaft region (r = –0.91, p < 0.0001), (Fig.2B.b), and Pearson’s correlation in the distal region (r = −0.61, p < 0.007), (Fig.2B.c).

### 3.3. Regional Micro-CT Assessment of Femoral Trabecular Structure

Three-dimensional reconstructions illustrate proximal metaphysis, midshaft cortex, and distal metaphysis for SCD-TMI-BMT and SCD-Con (Fig. 1B; panels d–k). As expected at the midshaft, where cortex predominates, no trabecular differences were detected, so trabecular analyses focus on proximal and distal regions.

At the proximal metaphysis, the bone surface to bone volume (BS/BV) ratio (Fig.2C. a) was higher in SCD-Con than SCD-TMI-BMT (43.70 vs 33.29 mm⁻¹, p=0.02; Fig. 2C. a), indicating a greater bone surface area relative to volume in SCD. This finding aligns with the higher trabecular thickness (Tb.Th) based on SCD-TMI-BMT (0.061 mm) compared to SCD (0.047 mm, p = 0.02, Fig.2C. b), suggesting a denser bone structure in the treated group. The structure model index (SMI), which describes trabecular morphology, differentiates between plate-like (close to 0) and rod-like (close to 3) structures. A higher SMI (2.29) in SCD indicates increased bone loss and fragility, whereas the lower SMI (1.34, p < 0.001) in SCD-TMI-BMT suggests a stronger, more load-bearing trabecular structure following treatment (Fig.2C. c). Additionally, the degree of anisotropy (DA), which measures the preferential orientation of trabeculae, indicates a more anisotropic trabecular structure in SCD (2.3) compared to a more isotropic structure in SCD-TMI-BMT (1.5), reflecting improved bone architecture in the treated group (Fig.2C. d).

At the distal metaphysics, Tb.Th remained higher in SCD-TMI-BMT (0.063mm) than SCD-Con (0.041 mm, p<0.05; Fig. 2B.e), and bone volume (BV) was greater in SCD-TMI-BMT (901.00 vs 809.73 mg HA/cm³, p=0.005; Fig. 2C.f). The complete set of micro-CT metrics (mean ± 95% CI) across all femoral regions for SCD-Con and SCD-TMI-BMT is provided in Supplementary Tables 1–3.

### 3.4. Histology

The H&E of femoral marrow (Fig. 2D) showed increased adipocyte spaces in SCD-TMI-BMT and diffuse hypercellularity in SCD-Con, consistent with erythroid hyperplasia. These qualitative findings paralleled higher PDFF and lower PDWF in the treated group.

## 4. Discussion

Quantitative MRI, complemented by micro-CT and histology, noninvasively delineated marrow and skeletal remodeling after total marrow irradiation with transplantation (TMI-BMT) in a transgenic SCD model. Across femoral regions, TMI-BMT mice showed lower PDWF, higher PDFF, and lower R2* than SCD controls, while micro-CT demonstrated thicker, more plate-like trabeculae with higher mineral density at metaphyseal sites; histology showed reduced cellularity with restoration of marrow adipocytes (larger/more adipocyte spaces). Together, these multimodal readouts indicate partial normalization of the marrow microenvironment and improved trabecular structure after marrow-selective conditioning(^6–9^).

### 4.1. Improved Water–Fat Balance in SCD mice after TMI-BMT

SCD marrow is classically hypercellular and fat-depleted owing to chronic erythropoietic drive and inflammation(^1, 2, 4^). Against that background, the coordinated decreased PDWF and increased PDFF after TMI-BMT across the proximal metaphysis, midshaft diaphysis, and distal metaphysis indicate a shift away from the water-rich, fat-poor SCD phenotype. Because PDWF and PDFF reflect the relative water and fat proton pools from chemical-shift separation, their reciprocal changes provide a sensitive, nonterminal readout of marrow composition(^15, 16^). In untreated SCD, persistent erythropoietic stress maintains a hypercellular, water-rich, fat-depleted marrow; the elevated PDWF we observed aligns with this inflammatory, hyperproliferative state, and reduced fat fractions are consistent with clinical and experimental reports of heightened hematopoietic activity in SCD(^15–17^). Quantitatively, group differences are shown for PDWF (Fig. 1C a–c) and PDFF (Fig. **1C** d–f), with representative parametric maps provided in Fig. 1D. Qualitatively, H&E sections (Fig. 2D) show reduced cellularity and enlarged adipocyte spaces after TMI-BMT, concordant with the PDFF increase.

### 4.2. Decreased Iron load after TMI-BMT

R2* (=1/T2*) was higher in SCD-Con and fell uniformly in TMI-BMT (main Group effect without Group×Region interaction), consistent with reduced susceptibility from iron/cellularity after treatment (Fig.1g–i; Fig. 2A c). Because paramagnetic iron and cellular packing dominate microscopic field inhomogeneities, *R2*\*↓ after TMI-BMT indicates a shift toward lower iron burden and more balanced composition (decreased PDWF /increased PDFF)(^2, 10, 12, 14^). As an independent biochemical surrogate of hemolysis and iron turnover, hepatic heme was lower in TMI-BMT than in SCD-Con, whereas serum heme did not differ (Fig. S2).

### 4.3. Skeletal and Architectural Considerations

Micro-CT demonstrated regionally consistent structural improvement after TMI-BMT. SCD-Con mice showed higher bone BS/BV and a rod-like, anisotropic network (higher structure model index [SMI], higher degree of anisotropy [DA]), indicating thinner, mechanically weaker trabeculae. In contrast, TMI-BMT mice exhibited greater trabecular thickness (Tb.Th), lower BS/BV, a shift in SMI toward plate-like architecture, and lower DA (more isotropic), features generally interpreted as mechanically favorable(^18, 19^) (Fig. 2C a–f). These structural findings align with clinical observations that poor trabecular quality contributes to fragility and low-impact fractures in SCD(^20^).

On the MRI side, susceptibility contrast was dominated by iron and cellular composition rather than trabecular interfaces. Although trabecular heterogeneity at 7 T can shorten T2* (raise R2*) by amplifying local field gradients, the between-group reduction in R2* was uniform across femoral regions, arguing that iron/fat changes outweighed architectural effects in this cohort(^21–23^) (Fig. 1C g–i). Mechanistically, iron overload (ferritin/hemosiderin) creates strong microscopic susceptibility gradients that accelerate spin dephasing (increased R2*), and, independent of imaging, adversely remodels bone by promoting osteoclastogenesis and suppressing osteoblast programs (lower Runx2 with reduced ALP/OCN and impaired mineralization)(^24, 25^). Thus, the SCD-Con pattern of thin, rod-like, anisotropic trabeculae is concordant with both higher iron burden and higher R2*, whereas the TMI-BMT shift toward thicker, plate-like, more isotropic architecture is directionally consistent with lower iron and lower R2*. Because conditioning and transplantation act in concert, changes in bone quantity/quality should be interpreted as the integrated outcome of TMI (myeloablation with organ sparing) and BMT (donor-driven reconstitution of the marrow niche), with MRI (Fig. 1C) and micro-CT (Fig. 2C) providing complementary evidence of skeletal remodeling.

### 4.4. Marrow Niche Rebalancing after TMI-BMT: Iron, MSCs, and Adipocytes

In SCD, chronic hemolysis and erythroid hyperdrive generate Iron overload which, together with inflammatory stress, impairs Mesenchymal Stem Cells (MSC) osteogenic/adipogenic differentiation, increases oxidative-stress–induced apoptosis, and weakens stromal support for hematopoiesis(^24–26^). SCD-derived MSCs also display intrinsic dysfunction via oxidative stress and TLR4/NF-κB activation, reducing HSC maintenance in coculture; functionally, donor HSCs cocultured with SCD MSCs engraft less well in myeloablated wild-type recipients than HSCs cocultured with wild-type MSCs, isolating a stromal contribution to engraftment failure(^26^). Notably, those assays used wild-type hosts and thus do not fully recapitulate the diseased SCD marrow milieu.

In our study, SCD mice received TMI conditioning followed by transplantation with AA (non-sickling) donor whole bone marrow. Post-TMI-BMT, mitigation of sickling/hemolysis plausibly lowers iron burden, creating conditions permissive for MSC recovery and lineage support. The direction of the imaging readouts, decreased PDWF/increased PDFF with lower R2* (Fig. 1C a-i), together with more plate-like metaphyseal architecture on micro-CT, (higher Tb.Th, lower SMI/DA; Fig. 2C a–f) is consistent with a niche transitioning away from hypercellular, iron-laden marrow toward partial adipose restoration (acknowledging that in vivo MSC function was not directly assayed).

Adipocyte effects are context-dependent. In physiologic wild-type marrow with intact stroma, expanded adiposity can oppose hematopoiesis/osteogenesis and hinder engraftment(^27^), whereas adipocyte-derived stem cell factor (SCF) can support HSC quiescence/regeneration during recovery(^28^). From the pathologically low baseline adiposity in SCD, a rise in PDFF after TMI-BMT likely reflects re-establishment of adipocyte–hematopoietic balance rather than maladaptive fat accrual, aligning with increased PDFF, decreased PDWF, and decreased R2*(^24–28^) (see Fig. 1C, 2C).

### 4.5. Integrative Interpretation and Clinical Implications

At four months post-TMI-BMT, the composite MRI signature, i.e., decreased R2* and decreased PDWF with increased PDFF, indicates decreased iron/cellularity and partial adipose restoration, directionally consistent with the micro-CT shift toward plate-like metaphyseal architecture (Fig. 2C a–c) and the uniform R2* group effect (Fig. 1C g–i). Clinically, PDFF/R2* are obtainable on MRI without contrast; vertebral/pelvic PDFF is reduced in SCD and tracks hemolysis severity, underscoring PDFF’s translational utility(^17^). Moreover, Iron overload in SCD has been associated with low bone mass, highlighting the value of monitoring marrow iron and bone quality alongside compositional MRI biomarkers(^29^). Taken together, our findings suggest a niche that is improved yet not fully regenerated at this time point, plausibly reflecting reduced iron load with suppression of erythropoiesis. The study was designed to isolate TMI-BMT-related changes and did not include wild-type comparators.

### 4.6. Complementary Roles of PDFF and R2* in Marrow Assessment

Across ROIs, marrow R2* and PDFF were inversely related, with a moderate-to-strong negative correlation when SCD-Con and SCD-TMI-BMT were pooled (Fig. 2B.a-c). The association was strongest at the mid-shaft, where trabeculae are sparse, supporting the view that in regions with minimal trabecular contribution, increases in adiposity further reduce local susceptibility variation and prolong T2* (i.e., lower R2*). Prior vertebral/femoral studies have similarly reported an inverse correlation between PDFF-R2*(^15, 16, 30^).

Because trabecular density/orientation, spatial iron deposition, and echo-time sampling can modulate decay, the relationship need not be strictly linear. In practice, the two metrics are complementary: PDFF captures the compositional shift from hematopoietic (water-rich) toward fatty balance, whereas R2* is more sensitive to susceptibility from iron and cellular packing. Consistent with this framework, the lower R2* observed in SCD-TMI-BMT aligns with reduced iron/cellularity, while higher R2* in SCD-Con reflects a water-rich, iron-laden microenvironment characteristic of ongoing erythropoietic stress.

### 4.7. Limitations and Future Directions

This study has four main limitations: (i) a modest, male-only cohort limiting generalizability; (ii) cross-sectional imaging at a single recovery time point, leaving trajectories of iron clearance, adipose restoration, and architectural change undefined; (iii) MRI susceptibility surrogates without quantitative iron assays and no ultrashort echo times (UTE), (TE <0.1 ms), reducing sensitivity to fast-decaying/bound-water components; and (iv) scale mismatch between MRI and micro-CT with ROI-level summaries, plus micro-CT/histology on a subset, limiting voxel-matched structure–function links.

Going forward, longitudinal imaging (pre-TMI, early post-conditioning, multiple recovery points), addition of UTE and quantitative susceptibility mapping (QSM), and region-matched voxel-wise MRI–micro-CT registration should refine attribution. Larger, sex-balanced cohorts and therapeutic studies (dose/fractionation, radiomitigators) can define parameters that optimize marrow/osseous recovery while preserving engraftment (9).

### 4.8. Conclusion

Multiparametric MRI and micro-CT delineated distinct marrow and bone profiles in SCD. Following TMI-BMT, marrow exhibited lower PDWF, higher PDFF, and lower R2*, alongside micro-CT evidence of thicker, more plate-like, and more isotropic trabeculae ( higher Tb.Th, lower SMI, lower DA, lower BS/BV), consistent with reduced iron burden and partial restoration of balanced marrow composition such as adipocyte, whereas SCD controls remained hypercellular and iron-rich. In this preclinical model, PDWF, PDFF, and R2* serve as sensitive, noninvasive compositional biomarkers of marrow remodeling, while micro-CT–derived metrics (Tb.Th, SMI, DA, BS/BV) provide complementary structural biomarkers of skeletal response. Together, these modalities support a practical framework for monitoring marrow health after marrow-selective conditioning in SCD.

## Acknowledgments

The authors would like to acknowledge the valuable contributions of the Small Animal Imaging Core (SAIC), Pathology Core, and Radiation Research Service Core (RRSC) at City of Hope National Medical Center.

## Author Contributions

Conceptualization, S.K.H.; Methodology, M.M., H.G., J.E.L., S.S.M. R.F, W.H, K.F, and, M.A.Z.; Formal Analysis, M.M.; Investigation, S.K.H. and S.S.M.; Writing-Original Draft Preparation, M.M.; Writing-Review And Editing, M.M., H.G., J.E.L, R.F., G.S., and S.K.H.; Visualization, M.M. and S.K.H.; Supervision, S.K.H; Project Administration, S.K.H.; Funding Acquisition, S.K.H. All authors have read and agreed to the published version of the manuscript.

