## Supplemental for "Quantitative MRI Reveals Bone Marrow Regeneration Following Targeted Marrow Irradiation and Transplantation in a Sickle Cell Disease Model"

**Supplementary Material**

1. **Mouse Model**

The mice were a well-established model of sickle cell disease incorporating targeted human globin gene modifications that were purchased from Jackson Laboratory, Maine, USA and housed at the COH facility during the study. Male Townes SS mice (CD45.2) B6;129-Hbbtm2(HBG1,HBB*)Tow/Hbbtm3(HBG1,HBB)Tow Hbatm1(HBA)Tow/J, sickling, aged 6 months, were used. Based on previous protocols with adjustments, the treatment plan for SS mice was modified to minimize lung and kidney exposure (by applying T spine beam isocenter) due to their radiosensitivity and risk of organ damage

The animal study protocol was approved by the Institutional Animal Care and Use Committee (IACUC) of City of Hope (COH) National Medical Center Duarte, CA, USA.

### 2. **MRI**

### 2.1.  **Mouse Positioning in MRI**

Anesthetized mice were placed in supine position in a body coil, and flat popsicle stick slabs were placed to align the femurs in a symmetrical manner. After securing supine position, a slight rightward rotation of the mouse helped achieve an improved coronal view, capturing the femur from proximal to distal end in a single imaging plane (Fig.1S.a). A body quadrature birdcage volume RF coil (38 mm × 70 mm) was used for all imaging, with gating applied as appropriate. Mice were anesthetized with 2–4% isoflurane in 2 L/min oxygen and maintained at 37 °C using a Minerve temperature control unit (temperature) set to 35 °C with real-time monitoring). Respiration (40–50 breaths/min) was monitored with an MR-compatible small animal gating system (SA Instruments, NY, USA). Localizers were acquired in the coronal plane to identify the position of the right femur. A water-filled tube was placed as a reference to monitor for any potential signal drift (Fig.1S.b).

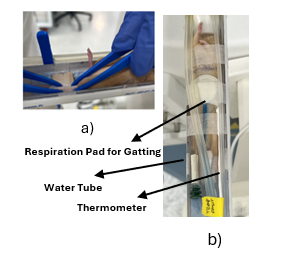

**Figure.1S. Mouse Positioning in MRI;** (a) Anesthetized mice were positioned supine in a body coil with femurs aligned using flat supports. (b) A respiratory gating attached to the thorax, a water-filled microtube attached to right femur for checking the signal drift correction and thermometer for monitoring the body temperature.

### 2.2. **MRI Acquisition**

B₀ and B₁ field homogeneities were well maintained, with B₀ variation within ±1 ppm over 20 mm along the z-axis, and B₁ variation within 0.05 axially (10 mm diameter) and 0.10 coronally (40 mm along z). B₀ was calculated using dual-echo GRE and B₁ using the VFA-AFI method^(1)^. For each mouse imaging session, automated B₀ shimming was performed prior to data acquisition to optimize magnetic field homogeneity. In addition to the standard auto-shimming protocol, further manual adjustments were applied.

2.3. **MRI Protocols**

A water-fat decomposition (6 echo times: 3.8–5.4 ms; TR = 200 ms; flip angle = 25°; FOV = 18×18 mm²; slice thickness/spacing = 0.5 mm; matrix = 128×128; NEX = 4; pixel size = 0.141×0.141 mm²) of the images was carried out using the globally optimal surface estimation algorithm (GOOSE), integrated into the preclinical scan software (v4.1.1.08) of the MR Solutions scanner. This algorithm effectively separates fat and water signals for accurate fat fraction quantification, with its primary advantage being the reduction of swap artifacts in fat-water decomposition^(2)^. The potential of PDFF MRI as a non-invasive, real-time in vivo biomarker for the quantitative assessment of bone marrow fat fraction in preclinical research was evaluated previously in our lab^(3)^.

For T2* mapping and accordingly R2* measurement, was conducted using fast low angle shot MRI (FLASH), spoiled multi echo gradient echo technique TR/TE 500ms/3, 5.52, and 8.04 ms. Voxel-wise signal decay was modeled using nonlinear least squares fitting (nlinfit) to the exponential decay function:

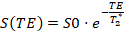
As

*S(TE)* is the signal at echo time TE, *S0* is the initial signal (amplitude at TE = 0).

2.4. **Region of Interest (ROI) Generation**

ROIs were manually delineated using the Image Segmenter application in MATLAB (MathWorks, Natick, MA). For each imaging modality, relevant slices were imported into the interface, and anatomical structures of interest were segmented using the polygon drawing tool, with careful attention to avoid the cortical bone and minimize partial volume effects. The resulting binary masks were saved as *.mat* files and utilized for quantitative analysis of PDFF/PDWF, and T2* relaxometry, as well as for calculating R2*. To reduce inter-operator variability, all ROIs were initially generated by a single experienced user; however, inter-operator variability was also assessed separately.

### 2.5.  **T2* mapping**

In these cases, the effective T2* values were often too short, and in some voxels, the decay occurred too rapidly to be reliably detected with the available echo times. This limitation reflects the inherent challenges of accurately quantifying T2* in highly inhomogeneous and fast-relaxing tissues at ultrahigh field strength. To address this, we implemented an upper threshold of 500 ms and a lower bound of 1 ms for fitted T2* values. This approach helped to account for voxel-wise variability and enhanced the reliability of comparisons across different marrow conditions. These imaging biomarkers provide important, non-invasive tools for monitoring bone marrow response in hematologic disorders and may help guide therapeutic decisions and post-treatment surveillance in SCD and other marrow-affecting conditions.

### 3. **Bone Preparation for micro-CT and H&E**

To enable sequential micro-CT and H&E while preserving mineral and cellular detail, right femurs of nine mice (4 SCD-Con/ 5 SCD-TMI) were excised (periosteum intact), fixed 24 h in formaldehyde/PBS, stored in PBS at 4 °C, stabilized for micro-CT, then decalcified for H&E. Routine formalin fixation and paraffin embedding for H&E staining dissolves lipids, especially unsaturated fats, due to their high solubility in ethanol and xylene, causing bone marrow adipocytes (BMAds) to appear as "ghosts" (empty spaces) that comes from fluid fat. This artifact distorts bone marrow adiposity (BMAT) assessment, particularly in aging and metabolic studies such as SCD mice, where saturated-to-unsaturated fat balance matters. We optimized this H&E staining preparation as follows:

#### 3.1. **Bone Marrow Histopathology**

The mice femurs were fixed in Z-fix (aqueous buffered zinc formalin fixative, ANATECH) for 2 days on a rocking platform, rinsed three times with Tris-buffered saline TBS (pH 7.4), and washed for 1 h in TBS. For decalcification, the samples were then incubated in freshly prepared 10% formic acid, with the solution replaced every hour for the first 3 hours, and then daily for two additional overnight incubations. Tissues were subsequently transferred to 10% formic acid for 2 days, followed by storage in 70% ethanol until they were processed for paraffin embedding and sectioning. They were then stained with Hematoxylin and Eosin using standard protocols. Whole-slide images of H&E-stained sections were captured using a digital slide scanner. Images were viewed and exported using **QuPath (v0.6.0)** for figure preparation and visualization^(4)^. Cellularity was evaluated on H&E-stained femoral sections by estimating the percentage of marrow space occupied by hematopoietic cells (myeloid, erythroid, and megakaryocytic elements) relative to adipose tissue and stromal space. Ten representative intertrabecular fields were examined at 10× magnification, and the percentage cellularity from each field was averaged to obtain a final value per sample^(5)^.

#### 3.2. **Heme Quantification Assay**

For sample preparation, 100 mg of Liver was homogenized in 100 µL of PBS, then centrifuged at 12,000 rpm for 10 minutes at 4 °C. The supernatant was collected and diluted 1:10 with distilled water. Total heme concentration was measured using a colorimetric Heme Assay Kit (ab272534; Abcam, Cambridge, UK) following the manufacturer’s protocol. The assay is based on the reaction of heme with a proprietary reagent under alkaline conditions to produce a colored complex measurable spectrophotometrically. All reagents and samples were equilibrated to room temperature (20–25 °C). For measurement, 50 µL of each sample was mixed with 200 µL of the Heme Assay Reagent in a 96-well microplate, alongside the prepared standards and distilled water as a blank. The plate was incubated for 5 minutes at room temperature, and the absorbance was recorded at 405 nm using a microplate reader. Heme concentrations were calculated from the standard curve using the following equation:

$$[\text{Heme}]=\frac{(\text{OD}_{\text{sample}}-\text{OD}_{\text{blank}})}{(\text{OD}_{\text{standard}}-\text{OD}_{\text{blank}})}\times C_{\text{standard}}\times D$$

where $C_{\text{standard}}$is the concentration of the standard (62.5 µM) and $D$is the dilution factor (e.i., 10) for each sample. All assays were performed in duplicate, and results were expressed as micromoles (µM) of heme^(6).^ The figure is shown in **Fig. S2**.

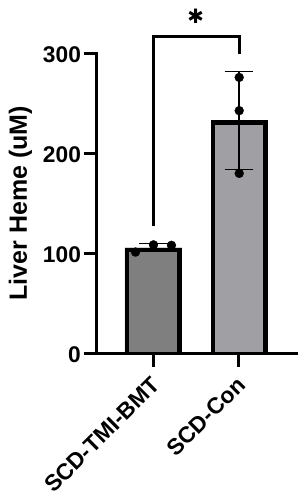

**Figure S2. Heme assay 15 weeks post-BMT.** (A) Liver heme (µM) was lower in TMI-BMT (106.5 ± 3.8, n = 3) versus SCD-Con (233.5 ± 44.7, n = 3); Mann–Whitney test, p = 0.011. Bars show mean ± SD; dots, individual mice.

**1. Micro-CT**

Femurs were imaged in wet gaze at 70 kVp (85 μA), 0.5 mm Aluminum filter, employing 720 cone beam projections per revolution and an integration time of 300 milliseconds within a field of view measuring 4.1 × 7.0 × 10.2 mm³. Three-dimensional images were reconstructed at 10 μm resolution using standard algorithms. Morphometric parameters were quantified directly and with the TRI model. Bone tissue mineral density was measured from micro-CT scans, calibrated using hydroxyapatite phantoms of known densities. The data were converted to true mineral density values (mg HA/cm³) using a standard linear attenuation curve, and the average density across threshold bone voxels in the volume of interest was reported by the Scanco dedicated software. Since our goal is to evaluate the effect of treatment on bone microstructure, we considered two analysis models. The Direct method is well-suited for diseased bone, while the TRI model, which assumes a plate-like trabecular structure, is more sensitive in healthy cases and early changes. Given that this study involves both disease progression and treatment response, we utilized both micro-CT analysis approaches to ensure comprehensive assessment. Morphometric parameters were included bone volume (BV), bone surface (BS), bone surface to bone volume (BS/BV), trabecular number (Tb.N), thickness (Tb.Th), spacing (Tb.Sp), connectivity density (Conn. D.), structure model index (SMI), mineral density values (mg HA/cm3), directional anisotropy (DA), and mid-shaft cortical bone mean density (Supplementary Table 1S, 2S, 3S).

**Table.1S.** Mean values with 95% confidence intervals of all extracted bone parameters analyzed across proximal site in SCD-Con and SCD-TMI-BMT groups.

| Parameter | Control (n = 4) (95% CI) | SCD-TMI_BMT (n = 5) (95% CI) |
| --- | --- | --- |
| Mean/Density [mg HA/ccm] - TV (Apparent) | 72.33 ± 75.38 | 105.28 ± 92.03 |
| Mean/Density [mg HA/ccm] - BV (Material) | 928.26 ± 70.55 | 946.64 ± 48.45 |
| TV [mm³] (Direct) | 0.78 ± 0.15 | 0.79 ± 0.13 |
| BV [mm³] (Direct) | 0.06 ± 0.05 | 0.11 ± 0.07 |
| BV/TV [1] (Direct) | 0.07 ± 0.05 | 0.14 ± 0.07 |
| Conn. D. [1/mm³] (Direct) | 27.47 ± 29.86 | 38.64 ± 20.63 |
| SMI [1] (Direct) | 2.28 ± 0.41 | 1.34 ± 0.54 |
| Tb.N* [1/mm] (Direct) | 3.66 ± 0.86 | 3.75 ± 0.30 |
| Tb.Th* [mm] (Direct) | 0.06 ± 0.02 | 0.08 ± 0.01 |
| Tb.Sp* [mm] (Direct) | 0.31 ± 0.08 | 0.29 ± 0.03 |
| TV [mm³] (TRI) | 0.77 ± 0.15 | 0.78 ± 0.12 |
| BV [mm³] (TRI) | 0.06 ± 0.05 | 0.11 ± 0.07 |
| BV/TV [1] (TRI) | 0.07 ± 0.05 | 0.13 ± 0.07 |
| BS [mm²] (TRI) | 2.45 ± 1.87 | 3.51 ± 2.10 |
| BS/BV [1/mm] (TRI) | 43.70 ± 11.46 | 33.29 ± 3.74 |
| Tb.N [1/mm] (TRI) | 1.54 ± 0.90 | 2.18 ± 0.98 |
| Tb.Th [mm] (TRI) | 0.05 ± 0.01 | 0.06 ± 0.01 |
| Tb.Sp [mm] (TRI) | 0.67 ± 0.40 | 0.46 ± 0.25 |
| [H1] [mm] | 0.59 ± 0.31 | 0 .45 ± 0.21 |
| [H2] [mm] | 0.98 ± 0.65 | 0.62 ± 0.30 |
| [H3] [mm] | 0.72 ± 0.38 | 0.52 ± 0.26 |
| DA [1] | 1.64 ± 0.34 | 1.38 ± 0.04 |

*Bone Volume (BV), Tissue Volume (TV), Bone Surface (BS), Trabecular Number (Tb.N), Thickness (Tb.Th), Spacing (Tb.Sp), Connectivity Density (Conn. D.), Structure Model Index (SMI), Mineral Density Values (Mg HA/Cm3), Directional Anisotropy (DA), Bone Surface To Bone Volume (BS/BV), Bone Volume (BV), Bone Volume Fraction (Bone Volume / Tissue Volume, (BV/TV), Connectivity Density (Conn. D), 3D image analyzer (TRI),*

**Table.2S.** Mean values with 95% confidence intervals of all extracted bone parameters analyzed across mid-shaft site in SCD-Con and SCD-TMI-BMT groups.

| Parameter | Control (n = 4) (95% CI) | SCD-TMI_BMT (n = 5) (95% CI) |
| --- | --- | --- |
| Mean/Density [mg HA/ccm] - TV (Apparent) | 200.51 ± 120.67 | 184.27 ± 97.24 |
| Mean/Density [mg HA/ccm] - BV (Material) | 1164.95 ± 56.00 | 1182.59 ± 28.50 |
| TV [mm³] (Direct) | 2.52 ± 0.77 | 2 .42 ± 0.61 |
| BV [mm³] (Direct) | 0.60 ± 0.31 | 0.56 ± 0.20 |
| BV/TV [1] (Direct) | 0.24 ± 0.10 | 0.23 ± 0.07 |
| Conn. D. [1/mm³] (Direct) | 1.52 ± 3.95 | 2.05 ± 3.99 |
| SMI [1] (Direct) | 1.49 ± 1.68 | 1.32 ± 2.36 |
| Tb.N* [1/mm] (Direct) | 5.98 ± 5.99 | 6.74 ± 3.08 |
| Tb.Th* [mm] (Direct) | 0.22 ± 0.11 | 0.21 ± 0.06 |
| Tb.Sp* [mm] (Direct) | 0.17 ± 0.27 | 0.07 ± 0.15 |
| TV [mm³] (TRI) | 2.50 ± 0.77 | 2.41 ± 0.61 |
| BV [mm³] (TRI) | 0.59 ± 0.30 | 0.55 ± 0.20 |
| BV/TV [1] (TRI) | 0.24 ± 0.10 | 0.23 ± 0.07 |
| BS [mm²] (TRI) | 5.43 ± 0.44 | 5.35 ± 1.09 |
| BS/BV [1/mm] (TRI) | 9.75 ± 3.99 | 9.99 ± 1.46 |
| Tb.N [1/mm] (TRI) | 1.11 ± 0.33 | 1.13 ± 0.21 |
| Tb.Th [mm] (TRI) | 0.22 ± 0.10 | 0.20 ± 0.03 |
| Tb.Sp [mm] (TRI) | 0.71 ± 0.24 | 0.70 ± 0.20 |
| [H1] [mm] | 0.67 ± 0.22 | 0.65 ± 0.12 |
| [H2] [mm] | 3.88 ± 2.21 | 4.59 ± 2.12 |
| [H3] [mm] | 0.83 ± 0.22 | 0.82 ± 0.17 |
| DA [1] | 5.82 ± 3.32 | 7.12 ± 3.24 |

*Bone Volume (BV), Tissue Volume (TV), Bone Surface (BS), Trabecular Number (Tb.N), Thickness (Tb.Th), Spacing (Tb.Sp), Connectivity Density (Conn. D.), Structure Model Index (SMI), Mineral Density Values (Mg HA/Cm3), Directional Anisotropy (DA), Bone Surface To Bone Volume (BS/BV), Bone Volume (BV), Bone Volume Fraction (Bone Volume / Tissue Volume, (BV/TV), Connectivity Density (Conn. D), 3D image analyzer (TRI).*

**Table.3S.** Mean values with 95% confidence intervals of all extracted bone parameters analyzed across distal site in SCD-Con and SCD-TMI-BMT groups.

| Parameter | Control (n = 4) (95% CI) | SCD-TMI_BMT (n = 5) (95% CI) |
| --- | --- | --- |
| Mean/Density [mg HA/ccm] - TV (Apparent) | 81.20 ± 115.10 | 46.39 ± 21.94 |
| Mean/Density [mg HA/ccm] - BV (Material) | 809.73 ± 21.00 | 901.02 ±52.35 |
| TV [mm³] (Direct) | 1.04 ± 0.81 | 1.33 ± 0.13 |
| BV [mm³] (Direct) | 0.09 ± 0.15 | 0.10 ± 0.03 |
| BV/TV [1] (Direct) | 0.07 ± 0.10 | 0.07 ± 0.02 |
| Conn. D. [1/mm³] (Direct) | 52.56 ±83.63 | 29.77 ± 16.30 |
| SMI [1] (Direct) | 2.99 ± 1.36 | 2.63 ± 0.32 |
| Tb.N* [1/mm] (Direct) | 4.14 ± 0.65 | 3.54 ± 0.37 |
| Tb.Th* [mm] (Direct) | 0.04 ± 0.02 | 0.06 ± 0.02 |
| Tb.Sp* [mm] (Direct) | 0.25 ± 0.03 | 0.30 ± 0.02 |
| TV [mm³] (TRI) | 1.03 ± 0.80 | 1.32 ± 0.13 |
| BV [mm³] (TRI) | 0.08 ± 0.14 | 0.09 ± 0.03 |
| BV/TV [1] (TRI) | 0.06 ± 0.10 | 0.07 ± 0.02 |
| BS [mm²] (TRI) | 4.34 ±5.99 | 3.98 ± 1.03 |
| BS/BV [1/mm] (TRI) | 71.35 ± 38.25 | 44.69 ± 10.48 |
| Tb.N [1/mm] (TRI) | 1.74 ± 1.94 | 1.51 ± 0.35 |
| Tb.Th [mm] (TRI) | 0.03 ± 0.02 | 0.05 ± 0.01 |
| Tb.Sp [mm] (TRI) | 1.01 ± 1.65 | 0.64 ± 0.18 |
| [H1] [mm] | 0.92 ± 1.41 | 0.59 ± 0.16 |
| [H2] [mm] | 1.18 ± 1.85 | 0.83 ± 0.23 |
| [H3] [mm] | 1.07 ± 1.74 | 0.68 ± 0.18 |
| DA [1] | 1.27 ± 0.13 | 1.40 ± 0.11 |

*Bone Volume (BV), Tissue Volume (TV), Bone Surface (BS), Trabecular Number (Tb.N), Thickness (Tb.Th), Spacing (Tb.Sp), Connectivity Density (Conn. D.), Structure Model Index (SMI), Mineral Density Values (Mg HA/Cm3), Directional Anisotropy (DA), Bone Surface To Bone Volume (BS/BV), Bone Volume (BV), Bone Volume Fraction (Bone Volume / Tissue Volume, (BV/TV), Connectivity Density (Conn. D), 3D image analyzer (TRI).*
